# spatzie: An R package for identifying significant transcription factor motif co-enrichment from enhancer-promoter interactions

**DOI:** 10.1101/2021.05.25.445606

**Authors:** Jennifer Hammelman, Konstantin Krismer, David K. Gifford

## Abstract

Genomic interactions provide important context to our understanding of the state of the genome. One question is whether specific transcription factor interactions give rise to genome organization. We introduce *spatzie*, an R package and a website that implements statistical tests for significant transcription factor motif cooperativity between enhancer-promoter interactions. We conducted controlled experiments under realistic simulated data from ChIP-seq to confirm spatzie is capable of discovering co-enriched motif interactions even in noisy conditions. We then use spatzie to investigate cell type specific transcription factor cooperativity within recent human ChIA-PET enhancer-promoter interaction data. The method is available online at https://spatzie.mit.edu.

## I. INTRODUCTION

Genome organization plays an important role in the function of the genome in development [1–3] and disease [4]. Specific transcription factor cooperation is a potential explanation for the cell type specificity of genomic interactions, especially those that tether enhancers to promoters. Recent methods seek to detect such transcription factor cooperativity by generating models to predict enhancer-promoter interactions and measuring the importance of model features [5,6]. However, these methods can be difficult to interpret either due to the complexity of model choice or the use of shrinkage techniques that could eliminate correlated features.

Here we introduce *spatzie*, an R/Bioconductor package named after the German diminutive for sparrow and inspired by the long-range geographical patterns of their songs [7], a reference to the long-range genomic interactions of transcription factor cooperativity. Within spatzie we implement a collection of statistical methods to identify transcription factor co-enrichment in experimental data obtained by protein-centric chromatin conformation methods such as ChIA-PET [8] and HiChIP [9]. We demonstrate the utility of spatzie by discovering the co-enrichment of transcription factor binding motifs simulated from ChIP-seq data. Furthermore, we apply spatzie to investigate cell type-specific interactions from RAD21-targeted ChIA-PET experiments across 24 human cell lines.

## II. MATERIALS AND METHODS

### A. ChIP-seq data for simulated co-enrichment

We simulate a cooperative relationship where binding of USF1 at promoters is co-dependent on binding of ELF1 at enhancers. Raw ChIP-seq data for USF1 and ELF1 from MEL mouse cells was downloaded from ENCODE (Supplementary Table S1). Reads were trimmed for adaptors and low-quality positions using Trimgalore (Cutadapt v0.6.2) [10] and aligned to the mouse genome (mm10) with bwa mem (v0.7.1.7) [11] with default parameters. Duplicates were removed with samtools (v1.7.2) [12] markdup, and ChIP binding events were called with GPS (v3.4) [13] with default parameters.

### B. ChIA-PET datasets

ChIA-PET interaction data was downloaded as processed files from Grubert et al. 2020 (Supplementary Material). Raw data for all experiments is accessible from GEO (Supplementary Table S1).

### C. Genomic annotations

For simulated data, mm10 promoter annotations were downloaded from the UCSC browser using the R package *GenomicFeatures* [14]. For human RAD21 ChIA-PET, hg19 promoter ensemble gene annotations were also downloaded from the UCSC browser using the same method. Within interaction data, regions that were within 2.5 kb of a promoter were classified as promoter regions. All other regions were classified as gene-distal enhancer regions.

### D. Statistical cooperativity calculation with spatzie

We implement three methods to measure the relationship between transcription factor binding motifs in promoter and enhancer regions of genomic interactions. Each method takes as input two vectors, ***x*** = (*x*_1_, *x*_2_, *. . ., x*_*n*−1_, *x*_*n*_) and ***y*** = (*y*_1_, *y*_2_, *. . ., y*_*n*−1_, *y*_*n*_), where *n* is the number of enhancer-promoter interactions. *x*_*i*_ is a set that contains all PWM scores for motif *a* in the promoter region of interaction *i*. *y*_*i*_, in contrast, contains those scores for motif *b* in the enhancer region of interaction *i*.

#### 1) Score-based correlation coefficient

We assume motif scores follow a normal distribution and are independent between enhancers and promoters. We can therefore compute how correlated scores of any two transcription factor motifs are between enhancer and promoter regions using Pearson’s product-moment correlation coefficient:

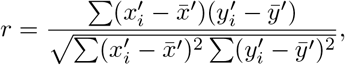

where the input vectors ***x*** and ***y*** from above are transformed to vectors ***x′*** and ***y′*** by replacing the set of scores with the maximum score for each region:

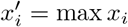

*x′*_*i*_ is then the maximum motif score of motif *a* in the promoter region of interaction *i*, *y′*_*i*_ is the maximum motif score of motif *b* in the enhancer region of interaction *i*, and 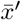 and 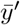 are the sample means.

Significance is then computed by transforming the correlation coefficient *r* to test statistic *t*, which is Student *t*-distributed with *n* − 2 degrees of freedom.

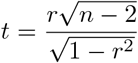

All p-values are calculated as one-tailed p-values of the probability that scores are greater than or equal to *r*.

#### 2) Count-based correlation coefficient

Instead of calculating the correlation of motif scores directly, the count-based correlation metric first tallies the number of instances of a given motif within an enhancer or a promoter region, which are defined as all positions in those regions with motif score p-values of less than 5 * 10^−5^, which tends to work well for human and mouse motifs [15,16]. Formally, the input vectors ***x*** and ***y*** are transformed to vectors ***x′′*** and ***y′′*** by replacing the set of scores with the cardinality of the set:

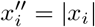

And analogous for *y′′*_*i*_. Finally, the correlation coefficient *r* between ***x′′*** and ***y′′*** and its associated significance are calculated as described above.

#### 3) Instance co-occurrence

Instance co-occurrence (or match association) uses the presence or absence of a motif within an enhancer or promoter to determine a statistically significant association, thus ***x′′′*** and ***y′′′*** are defined by:

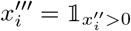

The significance of instance co-occurrence is determined by the hypergeometric test:

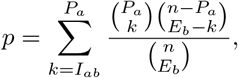

where *I*_*ab*_ is the number of interactions that contain a match for motif *a* in the promoter and motif *b* in the enhancer, *P*_*a*_ is the number of promoters that contain motif *a* 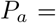 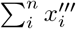, *E*_*b*_ is the number of enhancers that contain motif 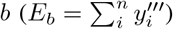, and *n* is the total number of interactions, which is equal to the number of promoters and to the number of enhancers.

#### 4) Multiple hypothesis testing

While the R package spatzie supports several methods to adjust p-values, three to control the family-wise error rate or FWER (Holm’s method [17], Hochberg’s method [18], Bonferroni’s method [19]) and two to control the false discovery rate or FDR (Benjamini and Hochberg’s method [20], and Benjamini and Yekutieli’s method [21]), all p-values presented in this work were corrected using the method of Benjamini and Hochberg.

## III. RESULTS

### A. spatzie tests transcription factor motifs for co-enrichment in enhancer-promoter interactions

The goal of spatzie is to identify pairs of transcription factor motifs which have a relationship such that the presence of motif A in an enhancer is associated with the presence of motif B in the promoter, indicating these transcription factors may be cooperating to drive enhancer-promoter interactions (Figure 1A). Given an input of interacting genomic loci, we select only those interactions which contain one locus that is gene-distal, which we label enhancer, and one locus overlapping a gene transcription start site, which we label promoter. Then, we scan these regions for transcription factor motifs using a database of DNA-binding motifs identified by ChIP-seq experiments, such as HOCOMOCO [22], HOMER [23], or JASPAR [24]. In order to limit hypothesis testing, spatzie provides a function to filter transcription factor motifs to those present in some threshold number of interactions. After filtering, we test transcription factor motifs pairwise for co-enrichment between enhancer and promoter pairs. Since the relationship between the DNA-binding motif and transcription factor activity is complex, spatzie provides three possibilities: 1) the strength of the transcription factor motif match (i.e., the PWM score), 2) the number of motif sites within the sequence, and 3) the presence or absence of motif sites. These definitions result in three different statistical tests for co-enrichment: the significance tests of 1) score-based or 2) count-based correlation coefficients, and 3) the hypergeometric test for co-occurrence over-representation (Figure 1B). Finally, we adjust the significance of these co-enrichment scores to account for multiple hypothesis testing and report significant transcription factor pairs.

**Fig. 1.**
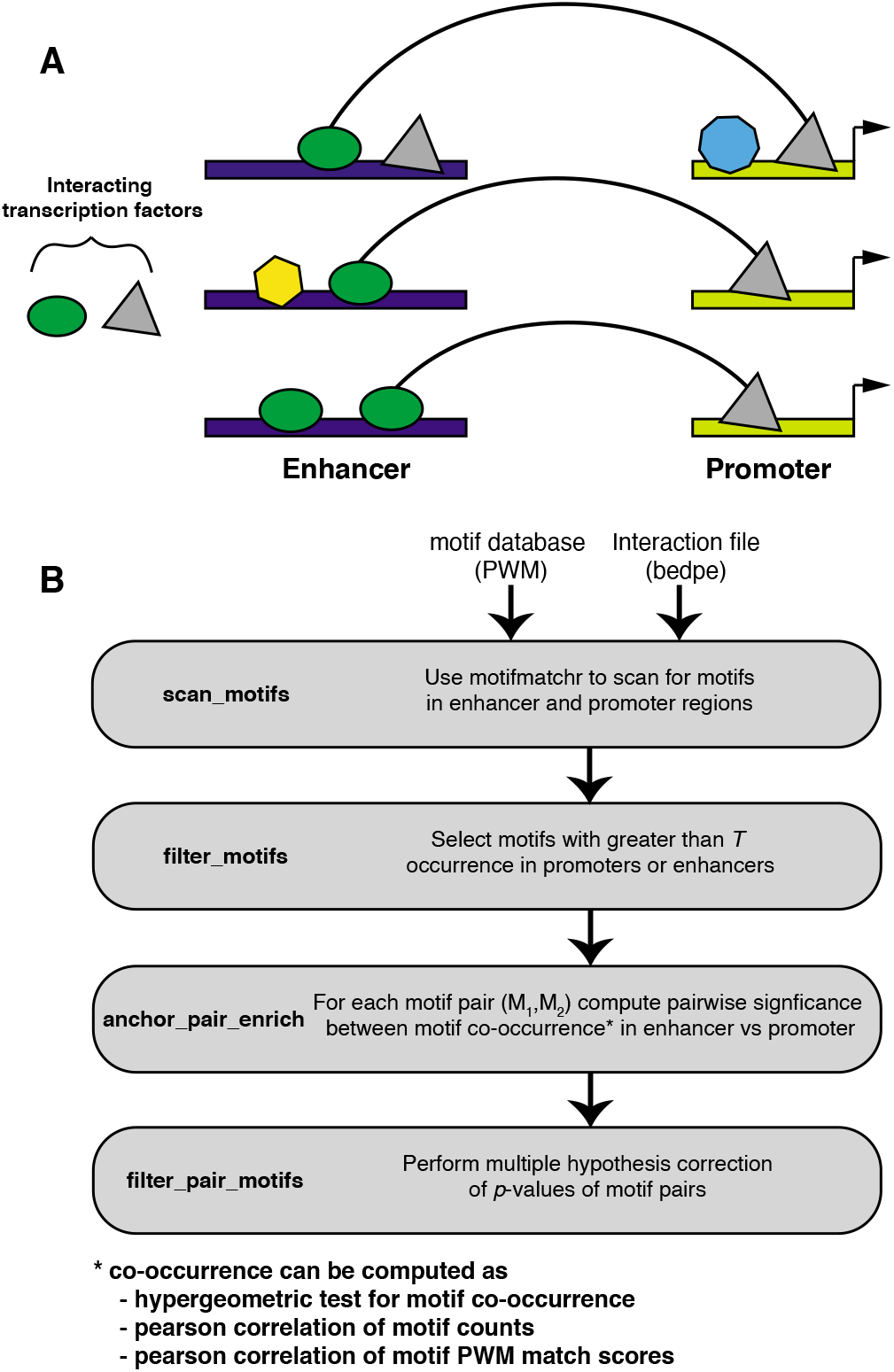
spatzie identifies motif pairs underlying enhancer-promoter interactions using co-occurrence and correlation statistics. **(A)** spatzie is designed to identify transcription factors which are facilitating interactions between enhancers and promoters based on detecting co-enrichment relationships between the presence of DNA-binding motifs in enhancer-promoter pairs. **(B)** Given input of a database of transcription factor DNA-binding motifs and a set of enhancer-promoter interactions, scan interactions for motifs, then limit analysis by filtering to motifs that are frequently present within the interactions of interest. Next we compute pairwise significance of motif co-occurrence in the enhancer and promoter data. Finally, we filter motif pairs that significantly co-occur under multiple hypothesis correction.

### B. spatzie identifies co-enrichment from simulated data

We validate spatzie by simulating a co-enrichment relationship where binding of USF1 at promoters is co-dependent on binding of ELF1 at enhancers. Using ELF1 and USF1 ChIP-seq data from ENCODE, we aligned and called binding events with GPS [13]. We then filtered USF1 binding sites to those that overlapped annotated promoters and filtered ELF1 binding events to any event that did not overlap a promoter. Then, we matched the most significant USF1 promoter event to the most significant ELF1 enhancer event, thus creating a simulated enhancer-promoter interaction data set where the strongest USF1 promoter events are matched with ELF1 enhancer events. We found that the three described methods (score-based correlation, count-based correlation, and motif presence/absence association) all result in significant co-occurrence between the USF1 motif and the ELF1 motif (Figure 2A-C). The DNA binding motifs of all other transcription factors with significant co-enrichment are highly similar to USF1 and ELF1 (Figure S1).

**Fig. 2.**
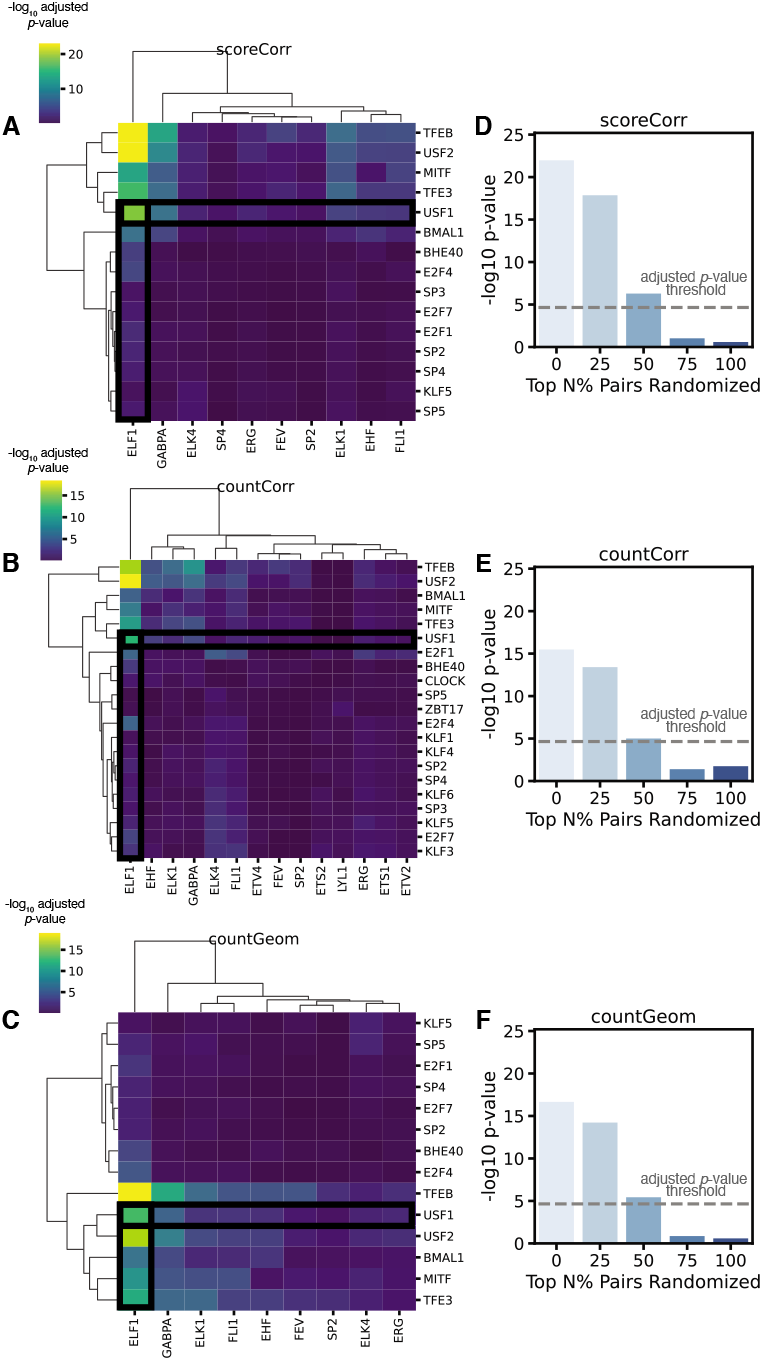
spatzie validates co-enrichment of ELF1 and USF1 on simulated enhancer-promoter interaction data. **(A)** spatzie cooperativity estimation computed using correlation of motif scores shows significant enhancer-promoter interactions for USF1 and ELF1 motifs. **(B)** spatzie cooperativity estimation computed using correlation of counts shows strongest enrichment between USF1 and ELF1 motifs. **(C)** spatzie cooperativity estimation computed using motif instance co-occurrence shows significant co-enrichment of USF2 and ELF1 motifs. Adjusted p-values were corrected with the Benjamini-Hochberg procedure. Randomization experiments where top *N* % of enhancer events are randomly permuted shows shrinking significance of co-enrichment under noisy data for **(D)** score correlation, **(E)** count correlation, and **(F)** hypergeometric co-enrichment. Dashed line represents the significance threshold at *p <* 0.05 under Bonferroni correction.

We also test that spatzie performs under noisy experimental conditions. We take the top *N* percent of enhancer interactions and randomly permute them such that they are paired with new promoters and then run spatzie using score-based correlation, count-based correlation, and motif presence/absence association (Figure 2D-F). We find that all methods collapse after 75% and 100% of the enhancers have been randomly permuted, indicating that co-occurrence is a result of the strength of the co-occurrence of the DNA binding motifs underlying the simulated enhancer-promoter interaction data.

### C. spatzie identifies germ layer and tissue-specific enhancer-promoter transcription factor interactions

Finally, we investigate enhancer-promoter interactions from RAD21 ChIA-PET of 24 human cell lines from ENCODE [25]. Based on the most significant co-enrichment scores on simulated data coming from the score correlation method, we chose to use score correlation to investigate transcription factor motif co-occurrence in enhancer-promoter interactions. After evaluating with spatzie the score correlation of 50,286 enhancer-promoter interactions that were present in at least one cell type, we find interactions cluster by tissue and germ line (Figure 3A-C), and that correlation of discovered motif pairs increases among cell types from the same germ layer and tissue (Figure 3D). While previous work has shown that cohesin-mediated genomic interactions are similarly stratified by germ layer and tissue [25], our analysis with spatzie shows that there is sufficient information within the co-enrichment of motifs underlying enhancer and promoter interactions to reproduce biologically meaningful germ layer and tissue layer organization. We found the tissue-level correlation between spatzie-discovered co-enriched motifs was reproducible with interaction calls using CID (Figure S2), which was previously shown to recover more reproducible interactions from ChIA-PET data [26]. We then extracted enhancer-promoter interactions that had the highest germ layer-specific expression and found that these include transcription factors such as Fox family members, which have a known role in endoderm development [27,28] and ectoderm development [29,30], and Nfatc4 in mesoderm interactions, which is a known T cell [31] and myogenic [32,33] differentiation factor. (Figure 3E). Similarly, we examined tissue-level specific interactions (Figure 3F) and found instances that were cell type specific, which is indicative of a potential trade-off in partnering of Sta5 at the enhancer with either Nfatc4 or Nanog at the promoter in blood or in liver tissues, respectively, or of AP2B at the promoter with Nfia or Zsc31 at the enhancers in breast or blood tissues, respectively (Figure 3F).

**Fig. 3.**
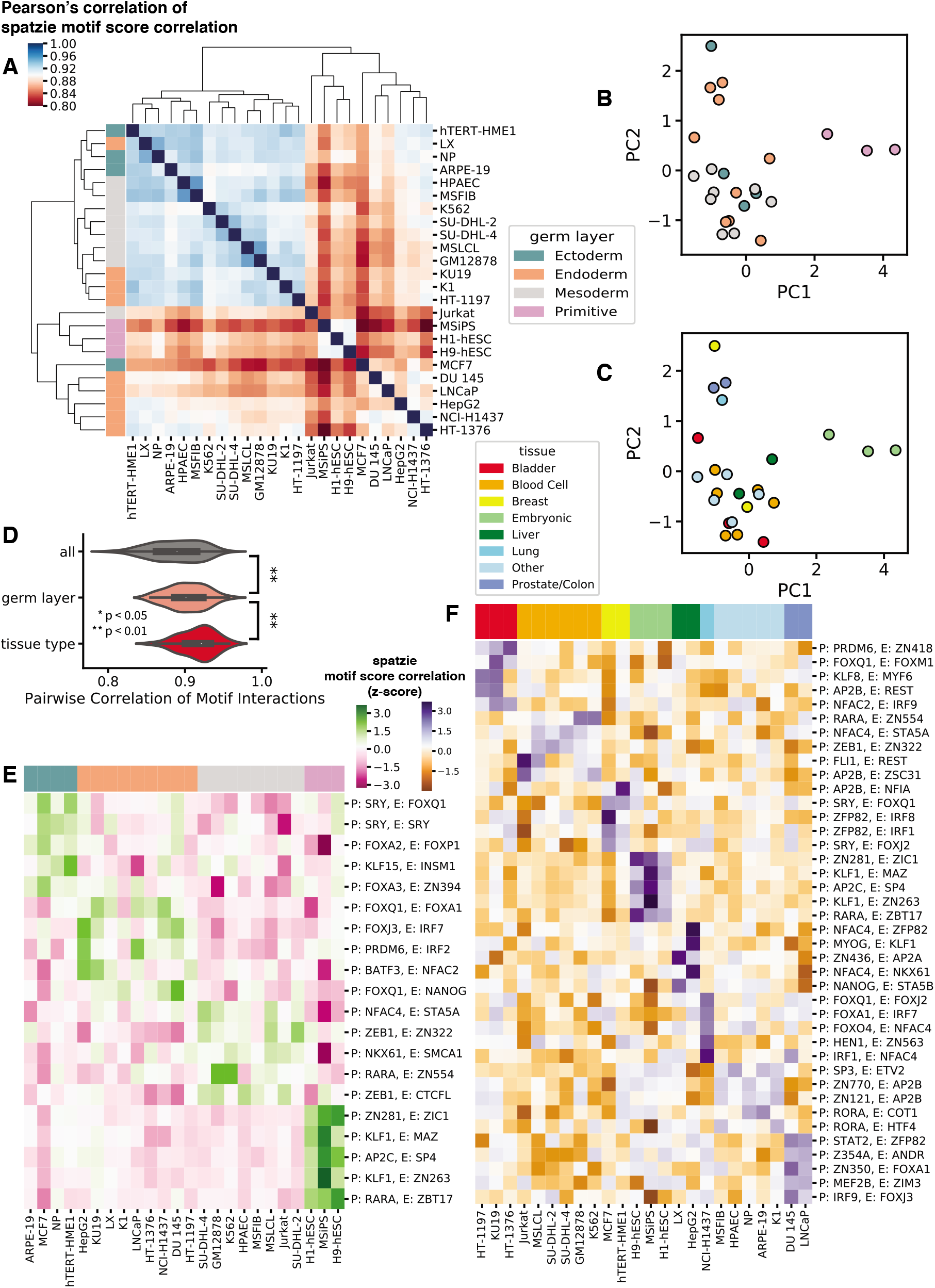
spatzie identifies transcription factor cooperation underlying interactions that are germ line and tissue-specific. **(A)** Pearson correlation of pairwise interactions shows similarity between related tissues. PCA on spatzie discovered transcription factor interactions shows **(B)** germ layer **(C)** and tissue type clustering. **(D)** Pairwise correlation of spatzie transcription factor motif interactions shows increasing relatedness of germ layer and tissue type. Significance computed by Wilcoxon rank-sum test. **(E)** Extraction of germ layer-specific transcription factor motif interactions include relevant lineage-determining transcription factor families, such as Fox in endoderm and ectoderm. Spatzie correlation scores are z-scores normalized by row. **(F)** Extraction of tissue-specific transcription factor motif interactions include potential transcription factor trade-offs at the promoter and enhancer that may mediate tissue-specific enhancer-promoter interactions. Spatzie correlation scores are z-scores normalized by row.

## IV. DISCUSSION

Overall, spatzie contributes to a growing field of tools for the analysis of enhancer-promoter interaction data by providing a collection of statistical tests to identify transcription factor motif co-enrichment. While other methods such as PEP-Motif [5] and the graphical lasso approach taken in Pliner et al. [6] may identify such co-enrichment relationships, they spend computational power to identify motifs that predict the activity of enhancers and promoters independent of their interactions, whereas spatzie focuses exclusively on identifying motifs which share a co-enrichment relationship between enhancer-promoter interactions. We validate spatzie on experimental data where we use real ChIP-seq data to simulate enhancer-promoter interactions between USF1 binding at promoters and ELF1 binding at enhancers. We show that spatzie’s three modes of motif pair relationship (motif score correlation, motif count correlation, and instance co-occurrence) all successfully identify USF1:ELF1 motif co-occurrence relationships even under noisy conditions, with motif score correlation achieving the most robust results. We also apply spatzie to data from 24 human cell lines and are able to show transcription factor co-enrichments that are discovered by spatzie cluster at germ-layer and tissue level, indicating these co-enrichment relationships are related to the organization of these cell types by germ layer and tissue. Furthermore, the identified germ layer and tissue-specific transcription factor interactions contain lineage-determining transcription factors, indicating that transcription factor co-enrichment between enhancers and promoters contains transcription factors that are known to play a role in differentiation and may point to their function as players in the structural organization of the genome. One concern with motif co-enrichment approaches is that the primary effects discovered can be attributed to the activity of cell type-specific transcription factors without a cooperative relationship. However, as evidence to the contrary we find examples of tissues that share enhancer or promoter motifs with different partners, indicating spatzie is identifying co-enrichment beyond general transcription factor activity. This combined with evidence that spatzie does not discover co-enrichment when we entirely randomize the relationship between binding for simulated interactions from USF1 and ELF1 ChIP-seq suggests that spatzie effects are not dominated by the general over-enrichment of motifs, but instead are based on a dependent relationship between a pair of transcription factors underlying enhancer-promoter interactions. In sum, we hope spatzie provides biological insight into the cell type-specific rules of transcription factor cooperativity underlying enhancer-promoter interactions.

## AVAILABILITY

The functionality of spatzie is bundled as an R package with the same name. The source code of the R package is hosted on GitHub (https://github.com/gifford-lab/spatzie). The core functionality is also available online at https://spatzie.mit.edu, which includes enhancer-promoter motif co-enrichment analysis with HOCOMOCO or user-defined motifs on interactions data mapped to either hg38, hg19, mm9, or mm10.

## AUTHOR CONTRIBUTIONS

Conceptualization, J.H.; Methodology, J.H. and K.K.; Software, J.H. and K.K.; Formal Analysis, J.H., K.K., and D.K.G.; Investigation, J.H. and K.K.; Resources, D.K.G.; Data Curation, J.H. and K.K.; Writing - Original Draft, J.H. and K.K.; Writing - Review & Editing, J.H., K.K., and D.K.G; Visualization, J.H.; Supervision, D.K.G.; Funding Acquisition, J.H. and D.K.G.

## ACKNOWLEDGEMENTS

We thank Michael Closser for helpful discussions in the conception of this project. We thank members of the Gifford and Wichterle labs for insightful suggestions and discussions.

