## Supplementary Material for "spatzie: An R package for identifying significant transcription factor motif co-enrichment from enhancer-promoter interactions"

### 1 Supplementary Figures

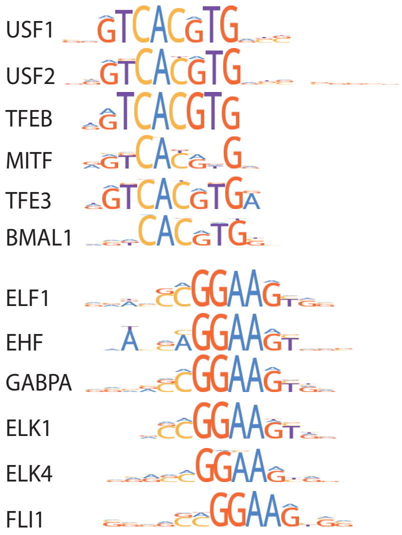


**Supplementary Figure S1:** Transcription factor motif position weight matrices that are discovered to have significant co-enrichment from USF1 and ELF1 simulated enhancer:promoter interaction task show high motif similarity to USF1 and ELF1 motifs.

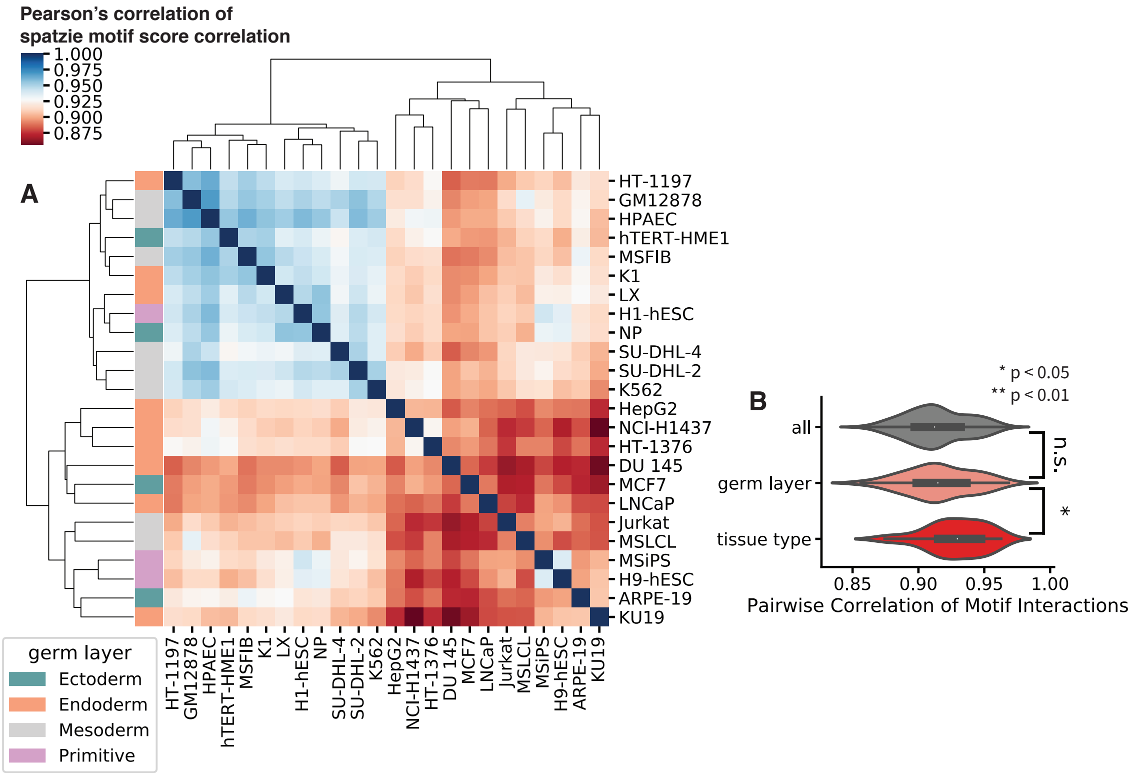


**Supplementary Figure S2:** **(A)** Pearson correlation of spatzie motif score of CID discovered enhancer:promoter interactions for RAD21 ChIA-PET of 24 human cell types. **(B)** Pairwise correlation of spatzie co-enriched motifs from CID enhancer:promoter interactions are higher among samples that are from the same tissue.

### 2 Supplementary Tables

| dataset label | Target | Cell line | GEO/ENCODE identifier |
|---|---|---|---|
| USF1 rep1 | ChIP | MEL | ENCFF996MWJ |
| USF1 rep2 | ChIP | MEL | ENCFF550MZG |
| ELF1 rep1 | ChIP | MEL | ENCFF186NAS |
| ELF1 rep2 | ChIP | MEL | ENCFF592ERV |
| ARPE | RAD21 | ARPE-19 | GSE134745 |
| DU145 | RAD21 | DU-145 | GSE134745 |
| MSFIB | RAD21 | fibroblast | GSE134745 |
| GM12878 | RAD21 | GM12878 | GSE134745 |
| H1 | RAD21 | H1 | GSE134745 |
| H9 | RAD21 | H9 | GSE134745 |
| LX | RAD21 | hepatocyte | GSE134745 |
| HepG2 | RAD21 | HepG2 | GSE134745 |
| HT1197 | RAD21 | HT-1197 | GSE134745 |
| HT1376 | RAD21 | HT-1376 | GSE134745 |
| HMTERT | RAD21 | hTERT-HME1 | GSE134745 |
| Jurkat | RAD21 | Jurkat-Clone-E6-1 | GSE134745 |
| AKTHY | RAD21 | K1 | GSE134745 |
| K562 | RAD21 | K562 | GSE134745 |
| KU19 | RAD21 | KU-19-19 | GSE134745 |
| LNCAP | RAD21 | LNCAP | GSE134745 |
| MCF7 | RAD21 | MCF-7 | GSE134745 |
| MSIPS | RAD21 | MSiPS | GSE134745 |
| MSLCL | RAD21 | MSLCL | GSE134745 |
| H1437 | RAD21 | NCI-H1437 | GSE134745 |
| NP | RAD21 | neural progenitor cells | GSE134745 |
| ECS | RAD21 | pulmonary artery endothelial cells | GSE134745 |
| DHL2 | RAD21 | SU-DHL-2 | GSE134745 |
| DHL4 | RAD21 | SU-DHL-4 | GSE134745 |

**Supplementary Table S1:** ChIP-seq and ChIA-PET datasets used in this study.

### 3 Supplementary Methods

For analysis of human RAD21 ChIA-PET data for co-enrichment of motifs underlying enhancer:promoter interactions, we applied spatzie to both the interactions that were provided by Grubert et al. using a custom method for generating a unified interaction set from many ChIA-PET samples and then using PET support to call cell type-specific interaction events as well as a unified interaction set that was generated by calling interactions with CID [1].

#### 3.1 Calling interaction events with CID

For each ChIA-PET experiment, we first used Mango 1.2.1 [3] (downloaded from `https://github.com/dphansti/mango`) to remove linker sequences and reads potentially due to polymerase chain reaction duplication, and aligned the raw reads using bowtie 1.2.3 (steps 1 - 3 in Mango pipeline). Mango was executed with the *reportallpairs* flag and the recommended parameter settings for the ChIA-PET Tn5 tagmentation protocol: *-keepempty TRUE -maxlength 1000 -shortreads FALSE*. CID (downloaded from https://groups.csail.mit.edu/cgs/gem/versions.html) was then run on the BEDPE file produced by Mango, using default parameters. Lastly, we used MICC [2] to assess the significance of all interactions identified by CID that are supported by more than one PET read.

##### 3.2 Generating a unified CID interaction set

For the CID interaction set, we first ran CID independently on all 48 samples (24 cell types; 2 replicates for each cell type). We then took the union of interactions called in all 48 samples. To merge overlapping interactions, we independently merged overlapping anchor1 and anchor2. We then removed interactions containing anchors that were larger than 20kb in size or interactions where as an artifact of merging, anchor1 and anchor2 overlapped. This resulted in a set of 71,643 joint CID interactions.

##### 3.3 Calling CID enhancer:promoter interactions for each cell type

We then obtained PET support from each of the 48 samples for the set of joint CID interactions and based on the correlation between replicates removed two samples (H1 rep2 and pulomonary artery endothelial cell rep2) that did not appear to have consistent correlation with their other cell type replicate and cell types within the same tissue. To obtain the interactions for each cell type from our joint CID interactions set, we looked at PET support and replicate correlation. For samples with two replicates, we called an interaction event within that cell type if it had greater than 1 PET in both replicates. For samples with one replicate, (H1 and pulomonary artery endothelial cell), we called an interaction event if the MICC FDR value of the event was less than 0.05. Interaction events were then filtered to those where one anchor was within 2.5kb of a promoter and the other was promoter distal (not within 2.5kb of a promoter). The selection criteria for calling cell type-specific allowed us to obtain a sufficient number of enhancer:promoter interaction events to run spatzie for co-motif enrichment (range 2,178-14,113 enhancer:promoter interactions per cell type) and were consistent with the range in the number of enhancer:promoter events discovered in Grubert et al. (range 1,476-11,850 enhancer:promoter interactions per cell type).

##### 3.4 Analysis of spatzie results for CID and Grubert et al. interactions

After co-enrichment of motifs at enhancer:promoter interactions was computed with spatzie, we analyzed the correlation between co-enrichment scores. For Grubert et al. interactions, we used only motif pairs where co-enrichment was significant under multiple hypothesis correction in at least one cell type. For CID interactions, we used all motif pairs as limiting to interactions that were significant under multiple hypothesis correction resulted in less meaningful clustering of motif co-enrichment scores by tissue type.
